# Diversification in the microlepidopteran genus *Mompha* (Lepidoptera: Gelechioidea: Momphidae) is explained more by tissue specificity than host plant family

**DOI:** 10.1101/466052

**Authors:** Daniel J. Bruzzese, David L. Wagner, Terry Harrison, Tania Jogesh, Rick P. Overson, Norman J. Wickett, Robert A. Raguso, Krissa A. Skogen

## Abstract

Insect herbivores and their hostplants constitute much of Earth’s described diversity, but how these often-specialized associations evolve and generate biodiversity is still not fully understood. We combined detailed hostplant data and comparative phylogenetic analyses of the lepidopteran family Momphidae to explore how shifts in the use of hostplant resources, not just hostplant taxonomy, contribute to the diversification of phytophagous insect lineages. We generated two phylogenies primarily from momphid species in the nominate genus, *Mompha* Hübner. A six-gene phylogeny was constructed with exemplars from Onagraceae hosts in western and southwestern USA and a cytochrome *c* oxidase subunit 1 (COI) phylogeny utilized both our collected sequences and publicly available accessions from the Barcode of Life Data System. Coalescent-based analyses combined with morphological data revealed ca. 56 undescribed *Mompha* species-level taxa, many of which are hostplant specialists on southwestern USA Onagraceae. Our phylogenetic reconstructions identified four major momphid clades: 1) an Onagraceae flower- and fruit-feeding clade, 2) a Melastomataceae galling clade, 3) an Onagraceae and Rubiaceae leafmining clade, and 4) a heterogeneous clade associated with multiple hostplant families, plant tissues, and larval feeding modes. Ancestral trait reconstructions on the COI tree identified leafmining on Onagraceae as the ancestral state for Momphidae. Cophylogenetic analyses detected loose phylogenetic tracking of hostplant taxa. Our study finds that shifts along three hostplant resource axes (hostplant taxon, plant tissue type, and larval feeding mode) contributed to the evolutionary success and diversification of *Mompha*. More transitions between exploited host tissue types than between hostplant families indicated that exploited host tissue (without a change in host) played an unexpectedly large role in the diversification of these moths.

## Introduction

Phytophagous insects comprise nearly a quarter of described metazoan diversity [1,2] and the majority of these insects specialize on a particular hostplant taxon [3,4]. Although the evolutionary processes behind these ubiquitous relationships have long garnered the attention of biologists [5,6], there is still much we do not understand about herbivore radiations. Colonization of novel hostplant taxa is hypothesized to be a dominant factor in the diversification of major phytophagous insect groups [7–10]. To the insect, new hostplant taxa serve as underexploited resources, and can provide enemy-free space or release from competition [11,12]. Once a phytophagous insect has shifted to a new hostplant taxon, local adaptation, assortative mating, and divergent selection can lead to isolation and ultimately speciation [13–17].

Shifts between hostplant species are well documented in phytophagous insect groups [18–23], but specific axes of the hostplant resource use are often overlooked. Diversification may also result from adaptation to a novel tissue type (e.g., leaves, fruits, or flowers) or a larval feeding mode (e.g., chewing, mining, galling, or boring) within the same host species [24,25]. Few previous studies have tracked patterns of colonization along multiple hostplant resource axes [14,26], and the relative lability, frequency, and dynamic nature of these shifts has not received appreciable quantification, especially in a fine-scale phylogenetic context. Here, we combine detailed hostplant data and phylogenetic comparative analyses to understand how shifts in the utilization of hostplant resources shaped the diversity of a phytophagous insect family that exploits many plant resource axes.

Momphidae (Lepidoptera: Gelechioidea) is a cosmopolitan family of small gelechioid microlepidopterans comprising approximately 120 named species [27]. Momphids are characterized by their small size and narrow forewings with raised scale tufts (Fig 1A). Recent studies of momphid genitalia and cytochrome *c* oxidase subunit 1 (COI) sequences indicate that the family is rich with cryptic species and undescribed taxa, warranting more intensive study of feeding niches, host plant usage, and systematic relationships across this group [28–31].

**Fig 1.**
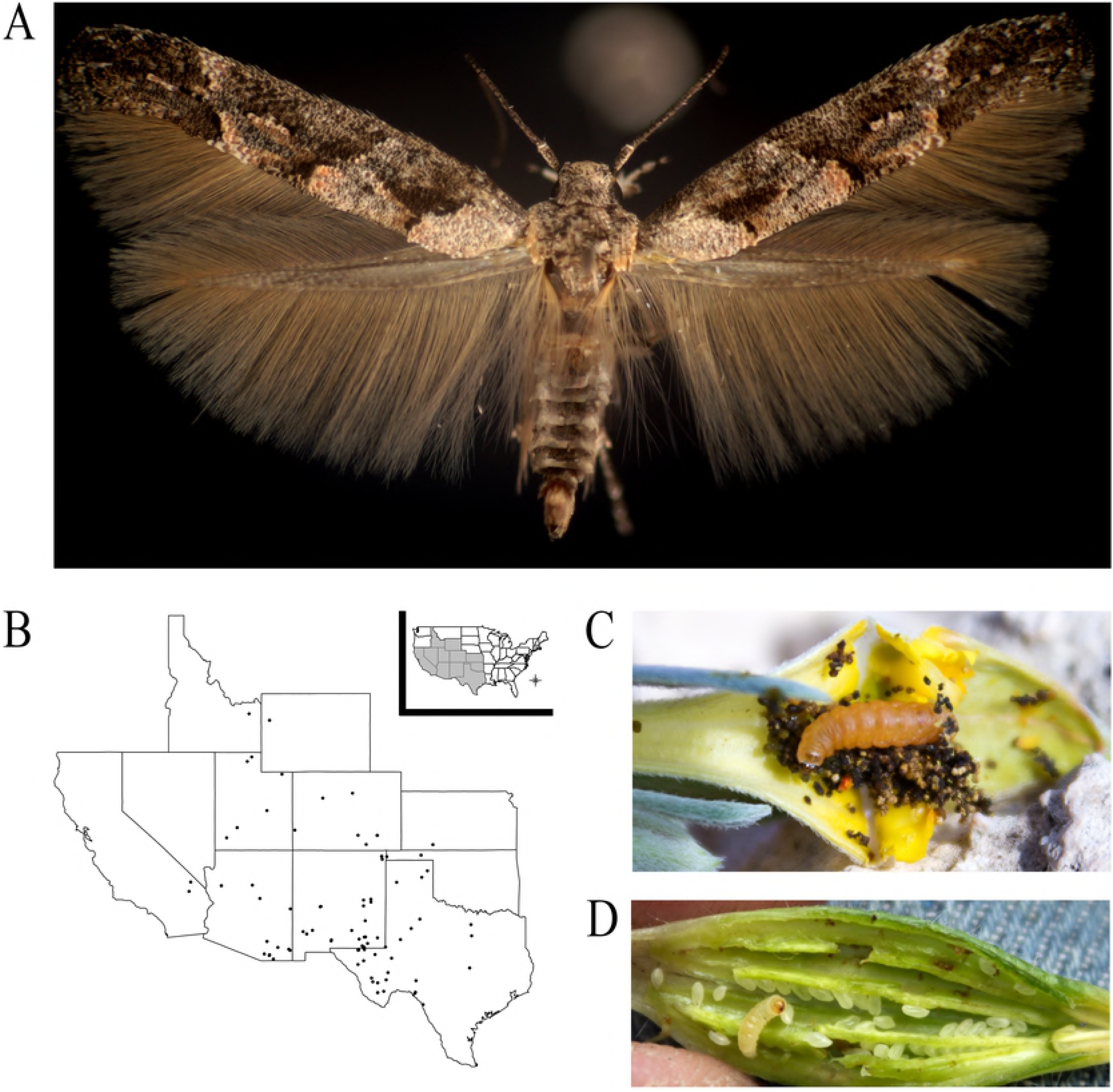
*Mompha* overview. A) *Mompha pecosella* Busck complex, Monahans Sandhills, TX from *Oenothera capillifolia* subspecies *berlandieri* W. L. Wagner & Hoch. Photo credit: T. Harrison B) Distribution of 87 *Mompha* collections from ten states: AZ, CA, CO, ID, KA, NM, OK, TX, UT, and WY. C) An opened *Oenothera lavandulifolia* Torr. & A. Gray flower bud showing a *Mompha* larva. Photo credit: R. P. Overson. D) An opened *Oenothera cespitosa* Nutt. fruit showing a *Mompha* larva. Photo credit: R. P. Overson.

Greater than 90% of described global momphid diversity is currently contained in the nominate genus *Mompha* Hübner (including *Zapyrastra* Meyrick, here nested within *Mompha*), whose larvae exploit a diverse array of hostplant resources. *Mompha* larvae mine, gall, and bore into six different plant tissues: flowers, fruits, leaves, shoot tips, stems, and roots. So far as is known, nearly all are monophagous or oligophagous hostplant specialists on one of seven dicot families: Onagraceae, Lythraceae, Rubiaceae, Polygonaceae, Cistaceae, Melastomataceae, and Haloragaceae [32–36]. Some *Mompha* also exhibit lability in hostplant tissue utilization between different broods [36] or instars [29]. More than 75% of *Mompha* species within North America are specialists on members of Onagraceae [28,29,36], suspected to have arisen from repeated radiations [35,37, this study].

In this study, we combine phylogenetic reconstruction and detailed hostplant-resource data (host taxonomy, plant tissue type, and larval feeding mode) to examine how shifts in hostplant resource axes contributed to the diversification of a species-rich insect group. We address the following questions: (1) What are the evolutionary relationships among *Mompha*? (2) What is the ancestral hostplant taxon, tissue resource, and feeding mode? (3) Are shifts to new hostplant tissues and changes in larval feeding modes as prevalent as shifts to new hostplant taxa? (4) Is there evidence of phylogenetic tracking between *Mompha* and their hostplants? We examined these questions at two scales: by reconstructing a six-gene phylogeny of Onagraceae-feeding *Mompha* for a subset of momphids from the southwestern USA, the center of diversity for Onagraceae and Nearctic momphid species [28,38] and by inferring a broad phylogeny of the family with COI barcodes.

## Materials and methods

### Sample collection

A total of 842 *Mompha* samples were included in the analyses. Of these, 131 were collected from 87 hostplant populations in the western and southwestern USA (Fig 1B) (S1 Table, MP accessions). Collections focused primarily on hostplants in *Oenothera* L., a large genus in Onagraceae with exceptional diversity in the southwestern USA. Larvae and pupae of *Mompha* were collected from flowers, fruits, leaves, shoot tips, stems, and roots when hostplants were at or near reproductive maturity (Fig 1C and 1D). Collected *Mompha* were fixed in 100% ethanol and stored at 40°C. Often, a second set of immatures was collected and reared to adulthood to aid in identification.

From the collected *Mompha*, we randomly selected one exemplar from each feeding resource axis: hostplant tissue type (flowers, fruits, leaves, shoot tips, stems, and roots), *Mompha* feeding mode (galling, boring, or mining), and hostplant family (Onagraceae, Lythraceae, Rubiaceae, Polygonaceae, Cistaceae, Melastomataceae, and Haloragaceae) within a hostplant population to sequence for five nuclear genes and COI. The accuracy of our representative sequencing scheme was verified by barcoding *Mompha* individuals with COI from all hostplants and all resource axes within ten hostplant populations. For each hostplant resource axis within each hostplant population, we found near-identical *Mompha* COI sequences, supporting the use of our representative sequencing scheme. An additional 47 adult *Mompha* were acquired for sequencing from the T. Harrison collection (Charleston, IL USA, S1 Table, TH accessions). To further assess taxonomic relationships, published COI barcodes for 664 *Mompha* accessions representing three continents and 15 countries (S1 Table, all other accessions) were downloaded from the Barcode of Life Data System (BOLD) (http://www.boldsystems.org) [39].

### DNA extraction, PCR, and sequencing

For the 178 previously unsequenced samples, DNA was extracted from either the anterior third of a caterpillar or a single adult leg, with a modified Chelex 100 and Proteinase-K protocol (S1 appendix) [40]. We amplified partial coding sequences from one mitochondrial locus (COI) and five nuclear loci [glyceraldehyde-3-phosphate dehydrogenase (GADPH), elongation factor 1-alpha (EF-1α), dopa decarboxylase (DDC), carbamoyl phosphate synthase domain protein (CAD), and histone 3 (H3)] (S2 Table). These loci have been used to reconstruct species-level relationships in other lepidopteran genera [41–44]. PCR was performed in 10 μL volumes (S1 appendix). Thermocycler programs were optimized for each primer pair (S1 Appendix). PCR product was verified with a 1% agarose gel stained with SYBR^®^ Safe DNA gel stain (Life Technologies, Grand Island, NY, USA) and stored at 4°C. For failed reactions, PCR was reattempted with internal primers generated for DDC, CAD, and GADPH (S2 Table). Successful amplicons were purified using Exonuclease I and Shrimp Alkaline Phosphatase (Affymetrix) (S1 Appendix). Purified PCR product was sequenced in the forward direction using a modified 10 μL BigDye^®^ Terminator v3.1 (ThermoFisher) cycle sequencing reaction (S1 Appendix) with standard thermocycler conditions (S1 Appendix). Sequenced product was purified with an EtOH/EDTA cleanup (S1 Appendix) and visualized on an ABI 3730 High-Throughput Sequencer.

Two datasets were generated for phylogenetic reconstruction: (1) *Mompha* collected in North America and sequenced for six genes (MP and TH *Mompha*), totaling 178 samples (six-gene dataset); (2) Pooled BOLD COI accessions and COI sequences from the six-gene dataset, totaling 842 individuals (COI dataset). Six outgroup species were selected from two recent gelechioid phylogenetic analyses [42,44]. Five of the six outgroup taxa fall within the scythridid assemblage: *Batrachedra praeangusta* (Haworth)*, Blastodacna hellerella* (Duponchel)*, Coleophora caelebipennella* Zeller*, Hieromantis kurokoi* Yasuda, and *Hypatopa binotella* Thunberg and one is from Gelechiidae: *Exoteleia dodecella* (L.).

### Phylogenetic analyses

For each locus, sequence chromatograms were converted to FASTA format in UGENE [45] and aligned at the nucleotide level with the FFT-NS-I algorithm in MAFFT v. 7.308 [46]. Aligned chromatograms were error-checked in parallel with the design tool in Genome Compiler (http://www.genomecompiler.com/). Low quality base calls were corrected with IUPAC nucleotide ambiguity codes in AliView [47]. AliView also verified open reading frames in amino acid alignments and trimmed sequences to equal length: CAD: 615 bp, COI: 558 bp, DDC: 288 bp, EF-1α: 474 bp, GADPH: 582 bp, H3: 276, bp. FASTA files were concatenated with SequenceMatrix v1.8 [48] to generate the six-gene dataset. Gene sequences were uploaded to GenBank (S1 Table).

Phylogenies were reconstructed with Maximum Likelihood (ML), Bayesian inference, and Coalescent methods using the CIPRES research computing resource [49]. Best-fitting substitution models and partitions for MrBayes and RAxML were selected with Bayesian Information Criterion (BIC) in PartitionFinder v1.1.1 [50] (S3 Table). For the six-gene and COI datasets, ML analyses were performed with RAxMLv8.2.9 [51] with 10 ML starting tree searches and 1000 bootstrap replicates using the inferred partitioning schemes and substitution models from PartitionFinder.

For both the six-gene and COI datasets, Bayesian analyses were carried out using MrBayes v3.2.6 [52]. For inferred partitioning schemes and substitution models, two independent runs were executed with three heated chains and one cold chain for 10 million generations sampled every 1000 generations. For the six-gene dataset, coalescent analysis was performed with four independent runs of StarBEAST2 [53], with bModelTest [54], unlinked strict estimated clocks for each locus, and default Yule priors for 150 million generations sampled every 5000 steps. For the COI dataset, BEAST2 v2.3.2 [55] was used to estimate relative divergence times of *Mompha* species. Four independent runs were executed with bModelTest, a strict molecular clock, and default Yule priors for 40 million generations sampled every 1000 steps. Convergence for each Bayesian and coalescent analysis was evaluated in TRACER [56] checking for ESS values greater than 200. If trees from an analysis converged, resulting trees were combined with a 10% burn-in and thinned to 10,000 trees with the program LogCombiner. Finally, a maximum clade credibility tree was generated for each analysis using median ancestor heights with the program TreeAnnotator. Agreement between tree topologies from ML, Bayesian, and Coalescent analyses for each dataset was evaluated using the APE [57] package in RStudio v3.3.1 [58] and FigTree v1.4.3 [59].

### Species delimitation

Morphological identification of *Mompha* exemplars was combined with three commonly used molecular species delimitation analyses to assign taxa to *Mompha* lineages in each dataset: (1) Discriminant Analysis of Principal Components (DAPC) [60], (2) Generalized Mixed Yule Coalescent (GMYC) [61], and (3) Poisson Tree Processes (PTP) [62]. DAPC is a multivariate analysis that uses aligned sequence data and the R package Adegenet [63]. GMYC is a likelihood method that compares branching events within and between species using an ultrametric tree and the SPLITS R package [64]. PTP is another likelihood method that compares the number of substitutions in branching events within and between species and is run with rooted phylogenetic trees. Three implementations were run: single rate PTP, Bayesian PTP, and Multi-Rate PTP [65]. Trees were trimmed to reflect delimited taxa.

### Character mapping

To investigate shifts in the utilization of *Mompha* hostplant resources, hostplant tissue type (flowers, fruits, leaves, shoot tips, stems, and roots), *Mompha* feeding mode (galling, boring, or mining), and hostplant family (Onagraceae, Lythraceae, Rubiaceae, Polygonaceae, Cistaceae, Melastomataceae, and Haloragaceae) were character coded onto the tips of the trimmed six-gene and COI *Mompha* phylogenies using GGTree [66]. Evolution of these discrete traits was traced with stochastic character mapping using 10,000 replicates with the R package Phytools [67].

### Cophylogenetic analyses

The cophylogeny of *Mompha* and its observed hostplants was examined at two scales: (1) *Mompha* with hostplants in Onagraceae, and (2) all *Mompha* and their recorded hostplant taxa. We used the host Onagraceae phylogeny (Overson et al., unpublished data) and for additional hostplant families, Phylomatic [68] generated a phylogeny of non-Onagraceae hosts that was grafted to the Onagraceae tree. For each analysis, an association matrix was generated that matched tips from the trimmed *Mompha* phylogeny to the tips of the host phylogeny. To evaluate if the *Mompha* phylogeny was non-randomly tracking the Onagraceae phylogeny, we used ParaFit [69] with 9,999 permutations to quantify the degree of congruence between host and insect topologies.

## Results

### Phylogenetic trees

The six-gene dataset consisted of a combined maximum of 2793 bp for 178 ingroup individuals: 150 (84%) of these were sequenced for all six markers and only one (~0.5%) individual was sequenced for fewer than four markers. ML and Bayesian phylogenies from the six-gene dataset, regardless of partitioning scheme, had congruent topologies, with robust support at shallow nodes separating *Mompha* clades. The coalescent phylogeny converged, but support for deep nodes was low (<0.75). Nevertheless, the topology from the coalescent phylogeny is largely consistent with the inferred ML and Bayesian phylogenies (10.5061/dryad.3n1g4td). Molecular delimitation analyses for the six-gene dataset recovered an estimated range of 21–37 *Mompha* species-level taxa (S4 Table): GMYC: 24 taxa; DPAC: 35 taxa; PTP: 30 taxa; bPTP: 37 taxa; and mPTP: 21 taxa. Morphological-based identification of *Mompha* exemplars combined with molecular delimitation, yielded approximately 31 *Mompha* species-level taxa (Fig 2). Seventeen species-level taxa were recognized as undescribed: of these, ten were collected in Central America and seven from the southwestern USA.

**Fig 2.**
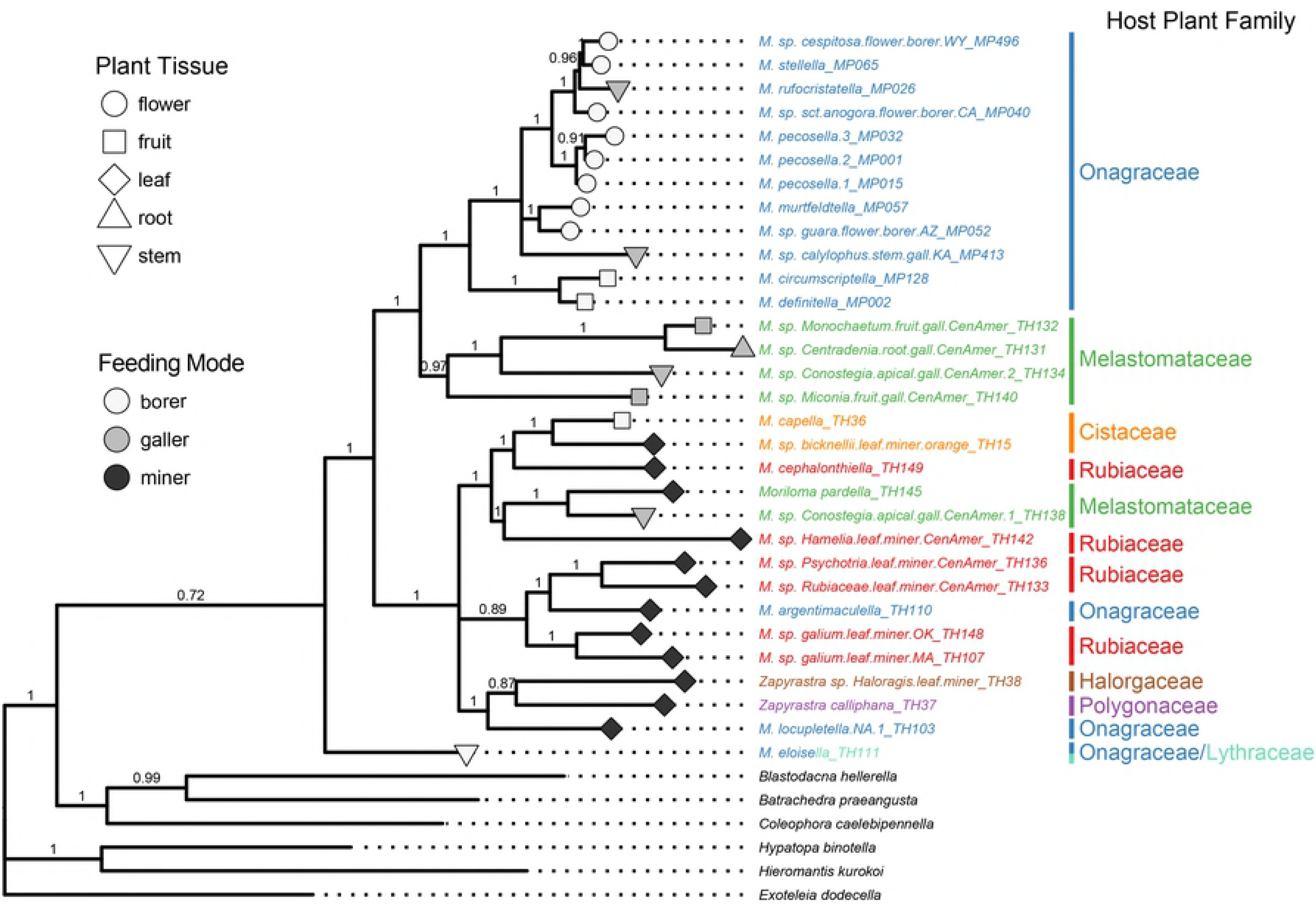
Bayesian phylogeny of the six-gene dataset. The phylogeny is trimmed to have one momphid species-level taxa per tip with posterior support shown at each node. When known, each tip was coded for hostplant family, hostplant tissue, and larval feeding mode. Four clades are shown: an Onagraceae-boring clade, a Melastomataceae-galling clade, and Onagraceae- and Rubiaceae-leafmining clade, and a heterogeneous clade that feeds on multiple hostplant families.

The combined COI dataset contained a maximum of 558 bp for 842 *Mompha* individuals. ML and Bayesian phylogenies had highly similar, but non-congruent topologies (10.5061/dryad.3n1g4td). The BEAST phylogeny was selected to represent the COI dataset because it is ultrametric and most similar to the reconstructed phylogeny of the six-gene dataset. As there were incongruences between the two datasets, we referred to the greater clade support in the six-gene dataset to resolve conflicts in topology. Molecular delimitation for the COI dataset recovered a range of 79–127 *Mompha* species-level taxa (S5 Table): GMYC: 87 taxa (CI: 84-95); DPCA: 86 taxa; PTP: 109 (p<0.001) taxa; bPTP: 127 (n=20,000) taxa; and mPTP: 79 taxa. Available morphological species identifications from exemplars in the six-gene dataset as well as distributional data were combined with molecular delimitation analyses to estimate 86 *Mompha* species-level taxa in the COI dataset. Of these, we find that possibly 56 represent undescribed taxa, most of which are endemic to northern latitudes, especially to southwestern USA (Fig 3).

**Fig 3.**
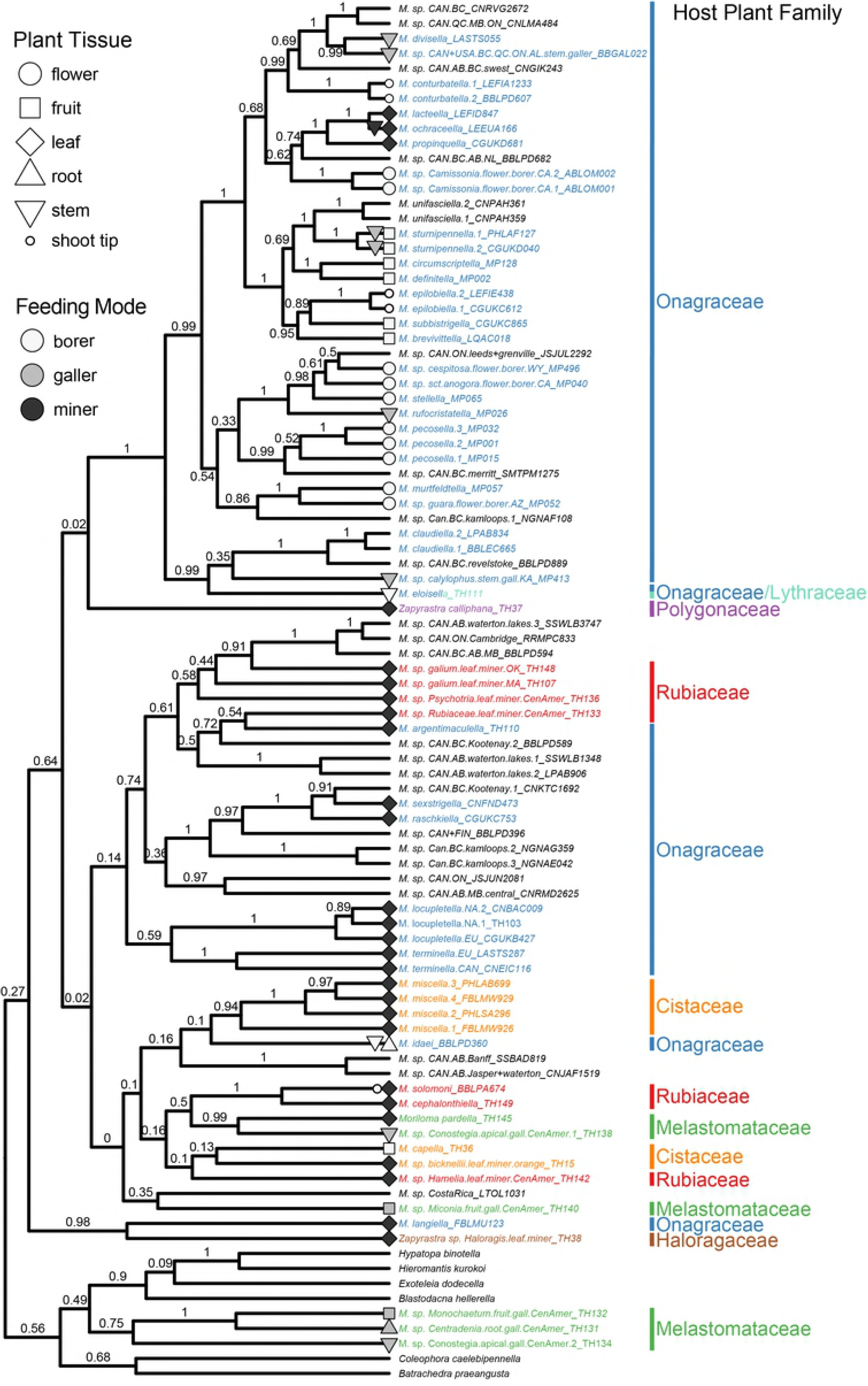
BEAST phylogeny of the COI dataset. The phylogeny is trimmed to have one momphid species-level taxon per tip. Posterior support is shown at each node. When known, each tip was coded for hostplant family, hostplant tissue, and larval feeding mode. Three clades are shown: an Onagraceae-boring clade, an Onagraceae- and Rubiaceae-leafmining clade, and a heterogeneous clade that feeds on multiple hostplant families. Here, the Melastomataceae-galling clade found in the six-gene phylogeny is split into a leafmining clade and a set of outgroup taxa.

### Ecological patterns and species relationships within *Mompha*

The six-gene phylogeny (Fig 2) supports four major clades of *Mompha*, each with a diagnostic host use pattern: (1) an Onagraceae-feeding clade that primarily bores into hostplant reproductive tissue; (2) a Melastomataceae-galling clade collected in Central America; (3) a leafmining clade that feeds on Onagraceae and Rubiaceae in both the Nearctic and Palearctic; and (4) a heterogeneous clade that feeds on multiple hostplant families. The COI phylogeny (Fig 3) recovers the same relationships as the six-gene phylogeny but with more taxa, except for the Melastomataceae-galling clade; some of its members cluster with outgroup taxa, while others nest within the leafmining clade.

### Character mapping

Ancestral trait reconstruction identified Onagraceae as the ancestral hostplant family (Fig 4). Shifts from Onagraceae to unknown were most common (S6 Table). The COI phylogeny identifies ten shifts to new hostplant families: one to Lythraceae, one to Polygonaceae, one to Haloragaceae, three to Rubiaceae, two to Melastomataceae, and two to Cistaceae. *Mompha* mine leaves on five hostplant families (Rubiaceae, Polygonaceae, Onagraceae, Haloragaceae, and Cistaceae), induce galls on plant tissue in two hostplant families (Onagraceae and Melastomataceae), and bore into plant tissue on three hostplant families (Onagraceae, Lythraceae, and Cistaceae). Galling and boring taxa have taxonomically restricted host associations, often feeding on a single hostplant species (see feeding records: S1 Table). Flower-boring arose twice (only in Onagraceae): once in *Oenothera* and once in *Camissonia* Link. Shoot-tip boring arose twice and only in Onagraceae. Fruit-boring arose twice, once each in Onagraceae and Cistaceae.

**Fig 4.**
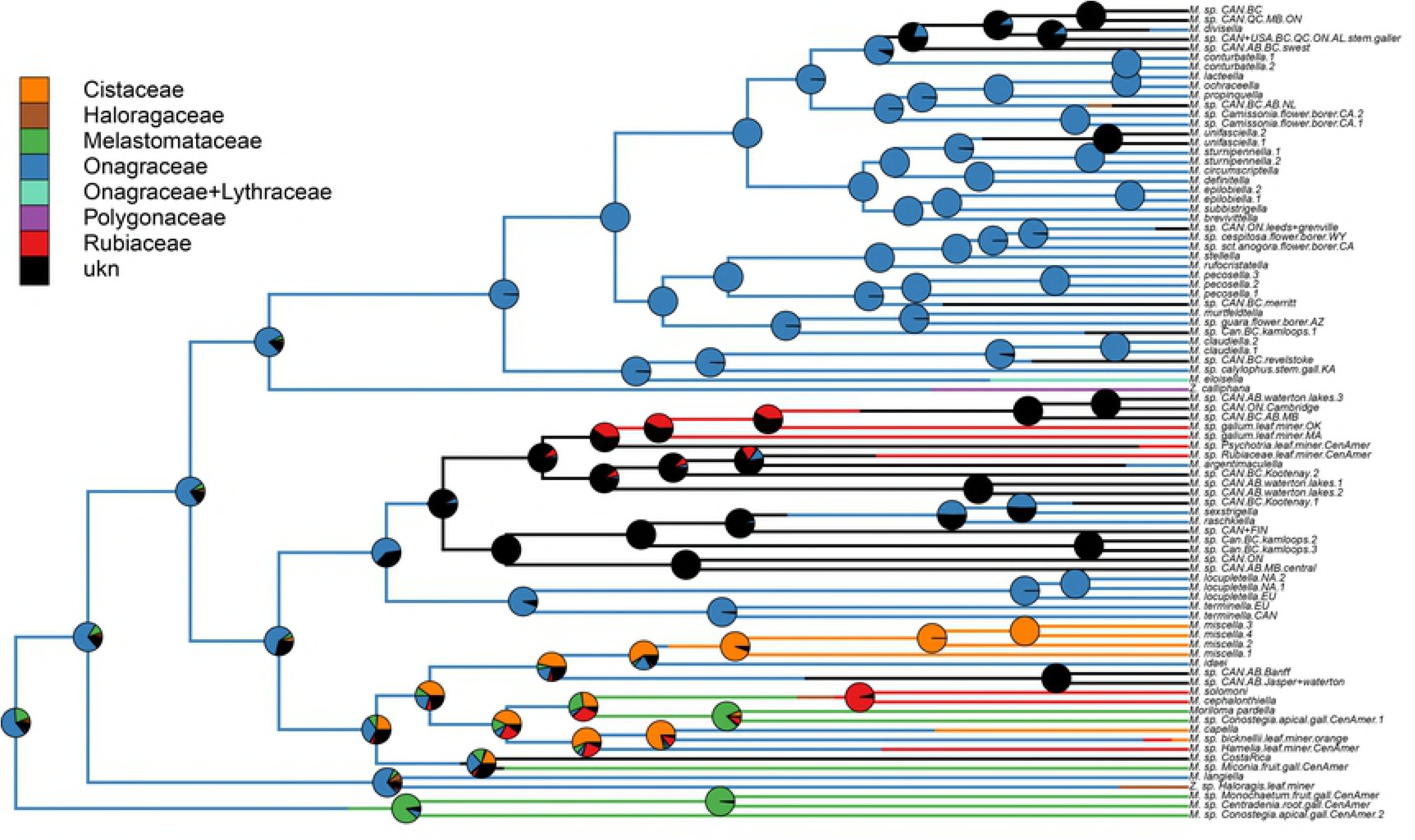
Ancestral trait reconstruction for *Mompha* hostplant family. Stochastic character mapping with 10,000 replicates. Posterior probabilities of ancestral states at each node are displayed as pie charts. Branch colors represent inferred character history.

Ancestral trait reconstruction identified leafmining as the ancestral utilized hostplant resource axis (Fig 5). Additional analyses that separate feeding mode and plant tissue type find that mining was the ancestral feeding mode and that leaves were the ancestral feeding tissue type (S1 and S2 Figs). The phylogeny identified 11 independent shifts from mining to new feeding modes. The most common shifts were from boring to unknown, mining to unknown, mining to boring, and from mining to galling (S7 Table). The COI phylogeny shows that *Mompha* shifted to new hostplant tissue in 17 instances. Shifts from leaf to unknown, from flower to unknown, and from leaf to stem + leaf were most prevalent (S8 Table). The Onagraceae-boring clade and the Melastomataceae-galling clade had the highest observed rates of switching between hostplant tissue types (Fig 3).

**Fig 5.**
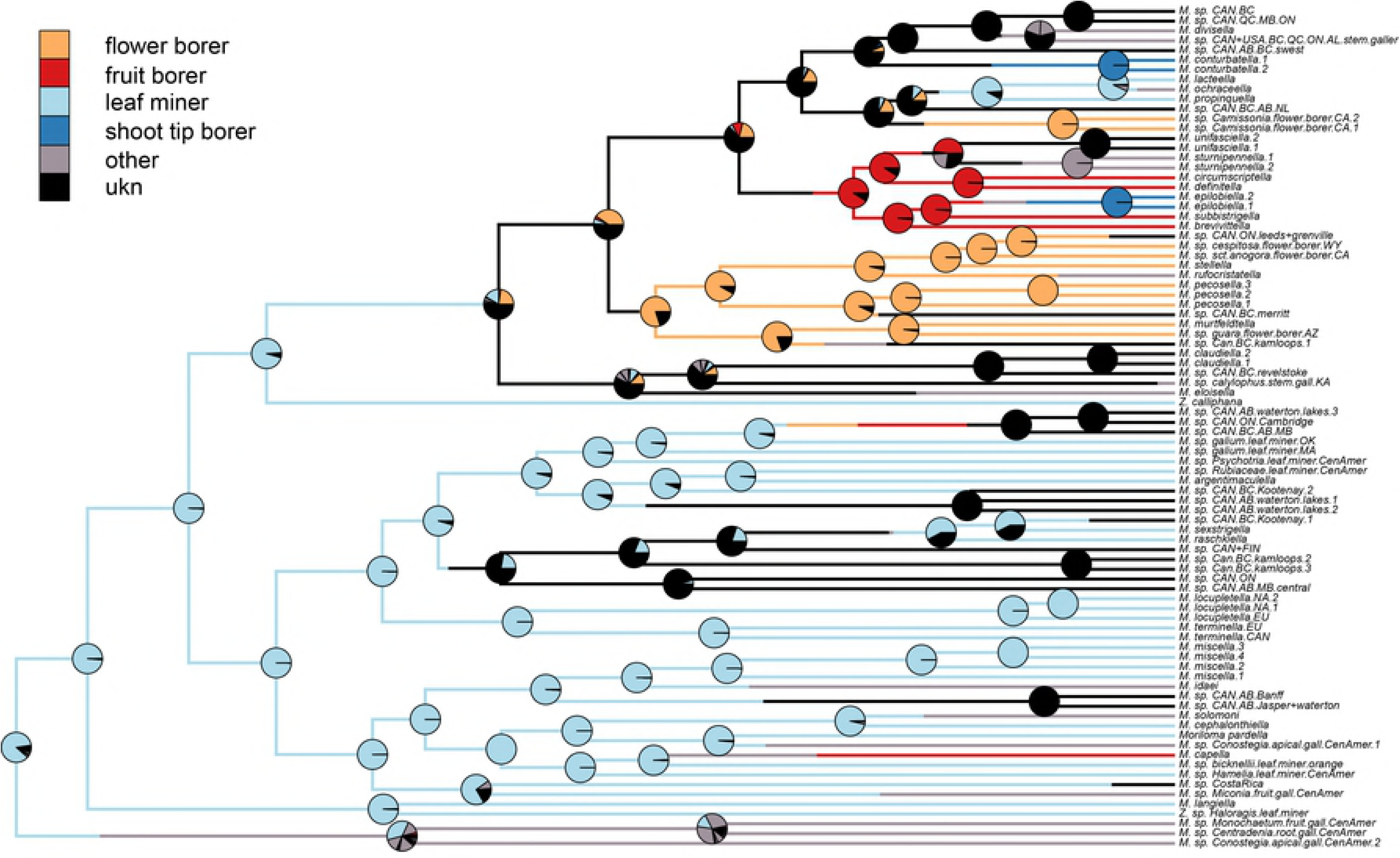
Ancestral trait reconstruction for *Mompha* hostplant resource. Stochastic character mapping with 10,000 replicates. Posterior probabilities of ancestral states at each node are displayed as pie charts. Both the branches and the pie charts are colored depending on the most likely character state at each node. Because of the many character states (15) found in the hostplant resource, we colored the most prevalent states and placed the remaining states in the “other” category.

Some *Mompha* taxa exhibited considerable lability in their use of hostplant resource axes. Four momphid taxa shifted hostplant tissue types: *Mompha sturnipennella* (Treitschke) galls stems in its first generation and bores into fruits in its second generation. In spring broods, early instars of *Mompha solomoni* Wagner, Adamski, and Brown are shoot tip borers but are only leafminers in their summer and fall broods. Early instars of *Mompha ochraceella* (Curtis) mine stems of their hosts, while later instars mine leaves. Finally, *Mompha idaei* (Zeller) bore into both the stems and roots of their hosts. *Mompha* taxa are also hostplant generalists, with some *Mompha* able to consume multiple hostplant species, sections, or sometimes genera (see feeding records, S1 Table).

### Cophylogenetic analyses

Distance-based cophylogenetic analyses found that *Mompha* loosely phylogenetically track the diversification patterns of their Onagraceae hosts (ParaFitGlobal = 3.46, p-value = <0.001) and the evolution of their known hostplants (ParaFitGlobal = 13.5, p-value = <0.001). While sister taxa of *Mompha* often shift to a closely related onagrad hostplant more frequently than a distantly related hostplant species, large jumps are not uncommon. For example, *Mompha rashkiella* (Zeller) and *Mompha sexstrigella* (Braun) both feed on *Chamaenerion* Ség. but their sister taxon, *Mompha argentimaculella* (Murtfeldt) feeds on multiple *Oenothera* species (Fig 6).

**Fig 6.**
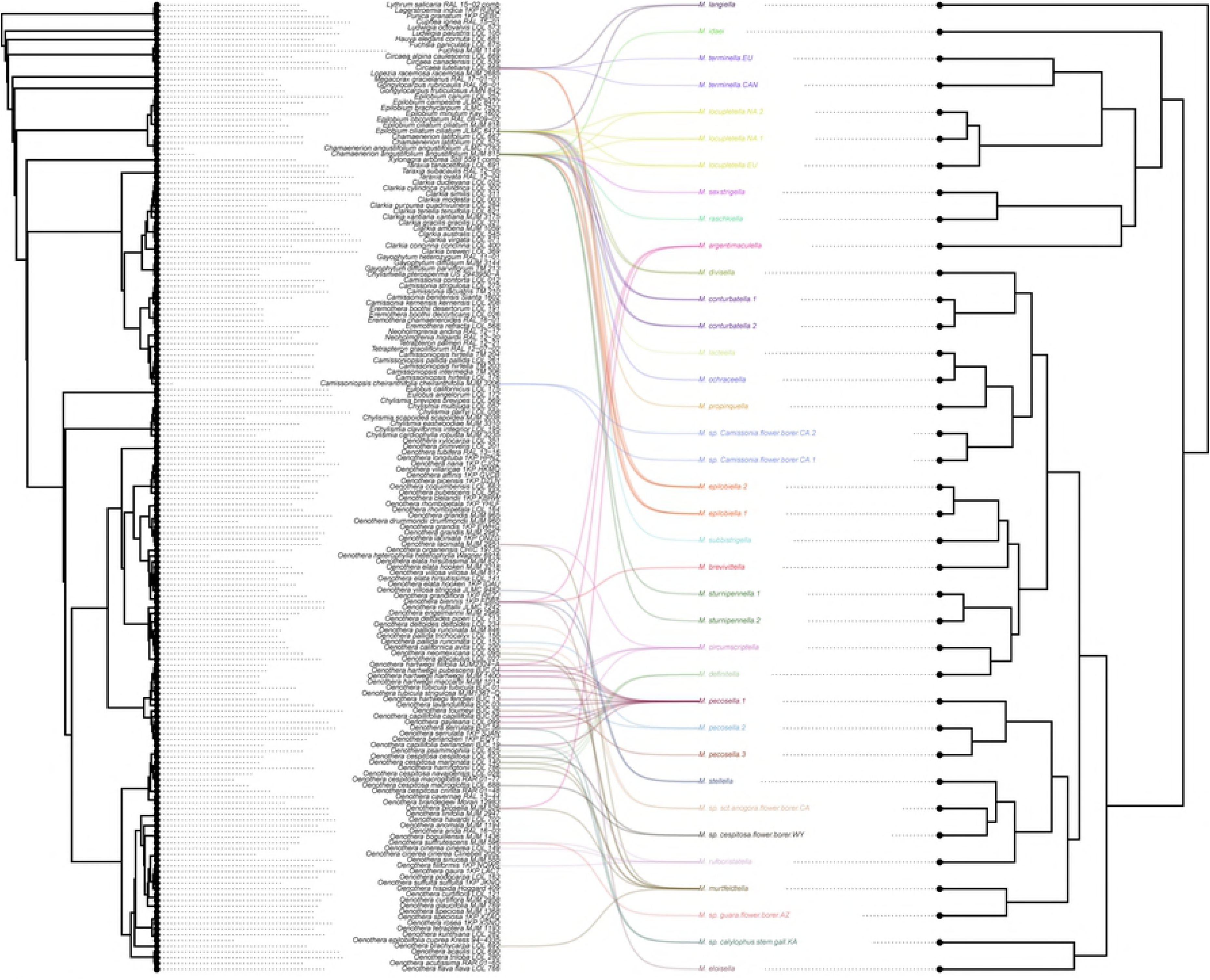
Aligned cophylogeny of *Mompha* and their Onagrad hosts. We used a Onagraceae phylogeny (Overson et al., unpublished data) and the COI *Mompha* phylogeny, trimmed to *Mompha* that use Onagraceae as their host. Linkages between *Mompha* and their hosts are colored for each species-level *Mompha* taxon. Tips of the *Mompha* phylogeny are rotated to minimize the distance between the host and Mompha taxa for visualization purposes.

## Discussion

We present two phylogenetic reconstructions of Momphidae (Gelechioidea) and document many new host records, life histories, and undescribed taxa in the nominate genus *Mompha.* Four well-supported clades were consistently recovered: an Onagraceae-flower and fruit-feeding clade, a Melastomataceae-galling clade, an Onagraceae- and Rubiaceae-leafmining clade, and a heterogeneous clade associated with many hostplant families and hostplant resources axes. We found that leafmining on Onagraceae was the ancestral feeding state and that *Mompha* shifted to new hostplant tissues more often than shifting to new hostplant families and larval feeding modes. The six-gene dataset supports the existence of 17 undescribed taxa (n=180 exemplars), while the larger COI dataset (n=848 exemplars) supports 56 undescribed taxa. The inclusion of Mompha from other hostplant taxa (beyond Onagraceae) and more extensive geographic representation may reveal additional undescribed taxa and provide greater insight into the evolutionary patterns of hostplant resource-axis switching than documented here.

### Host-mediated taxonomic diversification

Our examination of three hostplant resource axes suggests that *Mompha* diversification often coincides with shifts to new hostplant taxa, new plant tissue types, and / or changes in feeding modes. We infer that *Mompha* were ancestral leafminers on Onagraceae. From this ancestral state, *Mompha* more often shifted to a new plant tissue type, rather than to a new host family or feeding mode. Many of these shifts to new hostplant tissue types are unique to a few *Mompha* clades. For example, flower-boring only developed in the Onagraceae-flower and fruit-feeding clade, and appears to have evolved twice, once in *Oenothera*-feeding *Mompha* and then independently in *Camissonia*-feeding *Mompha*. Shoot-tip boring arose three times, twice in Onagraceae-feeding *Mompha* and once in Rubiaceae-feeding *Mompha*. Fruit-boring arose twice, once each in Onagraceae-feeding *Mompha* and the other instance in Cistaceae-feeding *Mompha*. While shifts to unrelated host taxa occur, species diversification across southwestern USA *Mompha* has been largely a matter of colonizing new hostplant congeners, especially within Onagraceae. It may be that internal feeding, where the host becomes both the food and the larva’s environment, leads to an increase in hostplant (lineage) fidelity in phytophagous insects [32,70,71].

Our data are relevant to the ongoing debate about the relative importance of the “musical chairs” hypothesis versus the “oscillation hypothesis” in generating herbivore species diversity [9,72,73]. Nearly all momphids are hostplant specialists, feeding on members of a single host genus or species group. It is more parsimonious to assume that the great diversity of this genus has come about through host switching within host lineages (e.g., within Onagraceae), with specialists begetting specialists (musical chairs hypothesis), than to assume that significant components of this radiation trace to dietary generalists that made host switches across lineages (oscillation hypothesis). A signal for the existence of dietary generalists in *Mompha* is lacking in our data. Host expansions (i.e., jumps to distantly related hostplant taxa) critical to the oscillation hypothesis, and important to generating taxonomic diversity in general, appear to have been modest within *Mompha*. Rather, the taxonomic diversification within the genus appears to be happening “in-house,” i.e., within the Onagraceae, through simple host, tissue, and feeding mode switching, fueled by the continuing radiation of their Onagraceae hosts.

Although we highlight the importance of shifts to new hostplant resource axes, we recognize that other evolutionary forces are also likely to have generated *Mompha* diversity. Both allochrony, and more prominently allopatry, without shifts in hostplant resources, contribute greatly to insect diversity [74,75]. The bud-boring *M. pecosella* Busck species complex and its hostplant ranges may provide an example. One member of the complex feeds exclusively on *Oenothera toumeyi* (Small) Tidestr, which is widely distributed in Sonora and Chihuahua with a few disjunct populations in the USA, found only above 1500 m in the Sky Island mountains of southeastern Arizona. A second *M. pecosella* lineage is only found in Utah and Colorado, even though its hostplants, *O. lavandulifolia* Torr. & A. Gray and *O. pallida* Lindl are broadly distributed across the southwestern and western USA. The third member of the complex feeds on several *Oenothera* species exclusively in Section *Calylophus* Spach across Texas, New Mexico, Arizona, and Oklahoma. Although the second and third element share *O. lavandulifolia* as hosts, they do not have overlapping ranges, suggesting allopatry played a role in the diversification of this species complex.

### *Mompha* and Onagraceae

Much of the known *Mompha* diversity is associated with Onagraceae, especially within *Oenothera*, *Epilobium* (L.), and *Chamaenerion*. *Mompha* and their onagrad hosts have formed a close evolutionary relationship in which *Mompha* sometimes impact hostplant fitness [76] and evolutionary trajectories of hosts at the population level [77,78]. *Oenothera*, particularly species in Sections *Calylophus* and *Pachylophus* Spach, host the majority of specialist-borer species of *Mompha* and are found in often arid areas. As in other phytophagous insect systems [74,79,80], plant phenology may be a driver of momphid diversification. Because many *Mompha* are strictly dependent on reproductive structures of *Oenothera* for larval survival, adult eclosion must be synchronized with the reproductive periods of their hostplants, which may be especially important in arid areas where precipitation and temperature modulate the timing of flowering [81,82].

Herbivore diversity often mirrors hostplant diversity [9,83,84]. The elevated alpha and beta species richness of *Oenothera* across the southwest USA, with ca. 75 species occurring within this region [85], coincided with higher species diversity of *Mompha*. Calibrated phylogenetic analyses of *Oenothera* and *Mompha* are needed to determine the relative timing and rates of their respective radiations, with the expectation that momphids, like other insect herbivores, will lag behind the diversification of their hosts [86].

In addition to onagrad taxonomic richness, *Mompha* niche partitioning on onagrad hostplants appears to have contributed to diversification across this genus. Our phylogenies document many instances of niche partitioning in *Mompha*, which allows for two or more species to share a hostplant. For example, we found three sympatric *Mompha* species on *Oenothera capillifolia* subspecies *berlandieri* occupying separate niches: stem-boring *M. rufocristatella* (Chambers), a flower-boring *M. pecosella* species, and an undescribed stem-galling species (*M. sp*. calylophus.stem.gall.KA). Niche partitioning was also documented in flower-boring *M. stellella* Busck and fruit-boring *M. brevivitella* (Clemens), which are sympatric and both use *Oenothera biennis* L. [87] at the same time of year. Similar niche differentiation is also seen in one of the best-studied model systems used to study co-diversification, the yucca moths (Prodoxidae) and their *Yucca* hosts. Most *Yucca* species host multiple species of *Prodoxus* Riley (stem and fruit borers) and *Tegeticula* Zeller (seed eaters) [88]. Thus, like the yucca moths, *Mompha* and their onagrad hostplants provide a promising system within which to study the effects of niche differentiation on diversification.

### Taxonomic considerations

*Mompha* is a monophyletic genus. However, our COI phylogeny and DAPC analysis identified *M. langiella* (Hübner) as sister to all other *Mompha* surveyed and placed it as a likely outgroup to the genus *Mompha*. Because *M. langiella* was only represented within the COI dataset, we refrain from formerly removing this species from the genus *Mompha*, until additional data can confirm its phylogenetic placement. In both datasets, species belonging to presumed outgroup genera nested within *Mompha*. For example, *Moriloma pardella* Busck, was placed as a member of the leafmining *Mompha* clade. The six-gene dataset suggests that the two *Zapyrastra* Meyrick species that were sampled in this study form a clade within *Mompha*; again, additional taxon sampling (including the type species *Z. calliphana* Meyrick) will be needed to support formal nomenclatural changes.

The limited morphological character differences among *Mompha* species and their small size make genetic barcoding a valuable source of taxonomic data [31,89]. The understudied alpha taxonomy of *Mompha* complicates the assignment of species names to our COI barcodes and underscores the need for taxonomic work on this genus. Barcoding revealed that several putatively Holarctic entities, in fact, represent separate, and sometimes multiple, Palearctic and Nearctic lineages [90,91]. In *Mompha*, we found that several Holarctic species complexes (e.g. *M. locupletella* (Denis & Schiffermüller) and *M. terminella* (Westwood)) should be separated into North American and European taxa. As was recently done with noctuid moths [92], collaborations between European and North American *Mompha* experts will help resolve species-level taxonomy of Holarctic taxa in this phenotypically challenging genus.

There remains many undescribed *Mompha* species in the Nearctic, Palearctic, and Neotropical regions. Considering only the Nearctic, where our study was anchored, there are 46 recognized *Mompha* species [93]. Our six-gene phylogeny suggest another seven undescribed species of *Mompha* north of Mexico, bringing the total to 53 potential Nearctic *Mompha* species. The average moth, microlepidopteran, and gelechioid genera in the Nearctic (north of the Mexican border) contain 5.2, 5.8, and 9.3 species, respectively [93]. By any of these measures, *Mompha* represents an extraordinarily speciose genus. Furthermore, the estimate of seven undescribed Nearctic species is likely conservative given that our COI phylogeny identified 33 additional undescribed species-level taxa, and that many Nearctic regions, plant communities, and hosts have yet to be sampled. In the Neotropics, tropical momphid diversity remains largely unstudied and it is highly likely that dozens of additional *Mompha* species remain undescribed. We have only identified six species level *Mompha* taxa that consume Melastomataceae, but melostomes are one of the most taxonomically rich and ecologically abundant Neotropical plant families, with over 150 genera [94] and thousands of species. The largest genus, *Miconia* Ruiz & Pav., contains 1050 estimated species [95] providing plenty of opportunity to contain undescribed *Mompha* diversity.

## Conclusion

We generated the first phylogenetic resources for *Mompha* and examined relationships between *Mompha* taxonomic diversity and three axes of hostplant resource usage. Our findings suggest that shifts to not only new hostplant taxa, but also host tissue types, and larval feeding modes, have been responsible for generating much taxonomic diversity across Momphidae. Our preliminary six-gene and COI phylogenies provide a framework to guide further collections and biosystematic studies of *Mompha.* We identified 56 undescribed *Mompha* taxa but given the geographically limited nature of our sampling, it is likely that many more *Mompha* await discovery. Future collection efforts should focus on the Melastomataceae of the Neotropics and Onagraceae in other areas globally, as our focus was on those in the western USA. Finally, calibrated reconstructions are needed to compare rates of diversification across the genus, map ecological changes, and better elucidate the role hostplants have in the diversification of *Mompha*, and thereby help to unravel and explain the exceptional diversity of this genus.

## Acknowledgments

The authors thank the following for assistance with fieldwork and sample collection: B. Cooper, M. Rhodes, E. Hilpman, E. Lewis, A. Gruver, L. Steger, and J. Fant. For providing valuable study material of *Mompha*, we thank M. J. Hatfield, G. Balogh, C. Eiseman, M. Palmer, R. Hoare, and B. Henning.

## Supporting Information Captions

**S1 Table**. Contains collection information for all *Mompha* contained in the phylogenies. Included are species names, collection location, hostplant taxa, hostplant resource, and GenBank ID. MP accessions denote southwestern collections. TH accessions denote Harrison collections. All other accessions taken from BOLD.

**S1 Appendix**. ***Mompha* sequence metadata.** Contains protocols used for DNA extraction, PCR, cleanup, and sequencing of samples.

**S2 Table**. **Primer sequence table.**

**S3 Table**. **Inferred partitioning schemes.** Generated with PartitionFinder using BIC model selection.

**S4 Table**. **Six-gene dataset combined species delimitation results**.

**S5 Table**. **COI dataset combined species delimitation results**.

**S6 Table. Mean *Mompha* state shifts for hostplant family**. Mean is taken from the 10,000 trees used for the ancestral trait reconstruction. Rows are the starting state and columns are the ending state.

**S7 Table. Mean *Mompha* state shifts for feeding mode**. Mean is taken from the 10,000 trees used for the ancestral trait reconstruction. Rows are the starting state and columns are the ending state.

**S8 Table. Mean *Mompha* state shifts for hostplant tissue**. Mean is taken from the 10,000 trees used for the ancestral trait reconstruction. Rows are the starting state and columns are the ending state.

**S1 Figure Ancestral trait reconstruction for *Mompha* larval feeding mode.** Stochastic character mapping with 10,000 replicates. Posterior probabilities at each node displayed as pie charts. Colors on branches and the pie charts represent most likely character state at each node.

**S2 Figure Ancestral trait reconstruction for hostplant tissue type.** Stochastic character mapping with 10,000 replicates. Posterior probabilities at each node displayed as pie charts. Colors on branches and the pie charts represent most likely character state at each node.

